# End-to-end Learning of Evolutionary Models to Find Coding Regions in Genome Alignments

**DOI:** 10.1101/2021.03.09.434414

**Authors:** Darvin Mertsch, Mario Stanke

## Abstract

1

**Motivation:** The comparison of genomes using models of molecular evolution is a powerful approach for finding or towards understanding functional elements. In particular, comparative genomics is a fundamental building brick in building high-quality, complete and consistent annotations of ever larger sets of alignable genomes.

**Results:** We here present our new program ClaMSA that classifies multiple sequence alignments using a phylogenetic model. It uses a novel continuous-time Markov chain machine learning layer, named CTMC, that is learned end-to-end together with (recurrent) neural networks for a learning task. We trained ClaMSA discriminately to classify aligned codon sequences that are candidates of coding regions into coding or non-coding and obtained six times fewer false positives for this task on vertebrate and fly alignments than existing methods at the same true positive rate. ClaMSA and the CTMC layer are general tools that could be used for other machine learning tasks on tree-related sequence data.

**Availability:** Freely from https://github.com/Gaius-Augustus/clamsa.

## 2 Introduction

Quantitative evolutionary models are important for many Bioinformatics tasks, such as phylogenetic tree construction or for testing hypotheses on whether an ORF is coding, for example in SARS-CoV-2 genomes [Jungreis et al., 2020]. Further, the data to leverage such evolutionary models from large numbers of genomes grows fast as evinced – for example — by the recent publication of alignments of the genomes of 240 mammalian species and 363 birds [Armstrong et al., 2020].

The evolution of biological sequences through substitutions is widely modeled with continuous-time Markov chains (CTMCs) on the state space of sequence characters and along the branches of a tree [Goldman and Yang, 1994, Lin et al., 2011, Ronquist et al., 2012, Stamatakis, 2014]. Commonly, the CTMCs are assumed to be stationary, homogeneous and time-reversible. Such a stochastic process is parameterized by a rate matrix alone that holds the rates of substitutions (state transitions) for any pair of different characters. Arguably most molecular phylogenetics programs estimate such a rate matrix *Q*, alongside the tree *T* that relates the sequences of an input multiple sequence alignment (MSA), e.g. the maximum-likelihood based program RAxML [Stamatakis, 2014] and the Bayesian method MrBayes [Ronquist et al., 2012].

Other applications of continuous-time Markov chains only require to estimate the rate matrix *for a given tree* and a multiple alignment of sequences observed at the leaves. Such applications are the identification of sites in a protein family that are under *positive selection*, e.g. with PAML [Yang et al., 1997], and the classification of genomic alignments into coding and non-coding, e.g. as done by PhyloCSF [Lin et al., 2011, Mudge et al., 2019]. Both of these widely used applications have in common that they require estimates of some parameterized form of a codon rate matrix that models evolution on the state space of 64 (or 61 sense) codons. For example, codeml from the PAML software [Yang et al., 1997] uses the codon substitution model GY94 of Goldman and Yang [Goldman and Yang, 1994] to estimate the rate ratio *ω* = *dN/dS* of synonymous and non-synonymous substitutions. The commonly used GY94 model has only few parameters besides *ω*. The estimated rate ratio ŵ has also been used for comparative gene prediction with AUGUSTUS-CGP [König et al., 2016].

We here consider the application to distinguish coding from non-coding regions in aligned genome sequences using CTMC models of codons. For this, Lin et al. train a so-called *empirical* general time-reversible model. Their program PhyloCSF uses rate matrices which are parameterized by the codon equilibrium distribution *π* and a triangular matrix totaling 63 + 63 ·64*/*2 = 2080 free parameters per rate matrix. Lin et al. trained two codon rate matrices *generatively* and independently using a maximum-likelihood criterion. One rate matrix models only coding regions and one rate matrix models only non-coding regions. These parameters are provided with PhyloCSF for a selection of animal clades as well as yeast but users cannot train PhyloCSF themselves. A comparison of the two respective likelihoods under two respective CTMCs is then used by PhyloCSF to classify codon or nucleotide multiple sequence alignments into coding or non-coding.

The selection metric *ω* can also be used for the same classification task, to distinguish coding and non-coding sequences, exploiting thereby the fact that, on average, most coding regions are under purifying (negative) selection and *ω <* 1. Such a *dN/dS test* that thresholds the estimate of *ω* was compared by Lin et al. to an earlier version of PhyloCSF and even performed best in their experiments [Lin et al., 2008]. PhyloCSF as introduced in Lin et al. [2011], however, performed somewhat better than the dN/dS test.

Siepel and Haussler [Siepel and Haussler, 2004] developed exoniPhy, which uses a hidden Markov model (HMM) of order 3 to segment a genome alignment into coding and non-coding regions. Its emission distribution is a CTMC model for an alignment column. exoniPhy also has three phylogenetic models for alignment columns in coding, non-coding and conserved non-coding genome regions.

A relatively high conservation of sequence is also an indicator that a region could be protein-coding [König et al., 2016]. However, such a predictive feature is prone to predict false positives in conserved non-coding regions. If only two CTMC models are used, one for coding and one for non-coding regions, then the different scale of their mutation rates is a nuisance parameter and atypical conservation hinders highly specific discrimination between the two. Lin et al. therefore *maximize* for each of the two models over a global mutation rate scaling factor in PhyloCSF. Siepel and Haussler coped with the problem by introducing a third model for conserved non-coding regions.

Recently, PhyloCSF has been used for a genome-wide scan to discover new human protein-coding genes, a “needle-in-a-haystack problem” [Mudge et al., 2019]. As PhyloCSF reports too many likely false potential coding regions, its output was further processed and filtered by a HMM and SVM whose parameters were trained while the ones from PhyloCSF were kept fixed. A relatively small candidate set of 658 approximate regions was then manually curated to annotate 70 previously undetected protein-coding genes. Although evolutionary models and comparative genomics are instrumental for identifying new coding regions that cannot be identified with other data, the high false positive number in a genome-wide scan remains a problem and the task to integrate the results into automatic gene structure annotation a challenge.

We here introduce a novel method for *end-to-end, discriminative* training of *arbitrarily many* rate matrices for a classification task. A new CTMC *layer* executes Felsenstein’s pruning algorithm in an automatically differentiable and therefore end-to-end trainable form. This layer can be combined with simple logistic regression, with neural networks or recurrent neural networks to evolutionarily classify multiple sequence alignments. Our program ClaMSA outperforms both the dN/dS test and PhyloCSF by a wide margin in the task of codon alignment classification. On vertebrate and *Drosophila* alignments ClaMSA predicts fewer than 16% of the false positives that PhyloCSF predicts at the same level of sensitivity. ClaMSA can be retrained by users for other tasks of evolutionary classification. However, a retraining for specific clades may not be necessary for a high performance.

ClaMSA collects possibly powerful evolutionary evidence that a candidate exon or gene is indeed protein-coding and conserved. The versatile CTMC layer module can be used independently for other machine learning tasks that learn continuous-time Markov chains on trees. The code for both is freely available under https://github.com/liemonade/clamsa.

## 3 Methods

We first present the new CTMC layer for evolutionary data and then how our new program ClaMSA uses mostly established machine learning components and methods to classify multiple sequence alignments.

### 3.1 CTMC Layer

Our model is the standard *general-time reversible* (GTR) CTMC on a tree that we recall in the Sup-plementary Data for convenience. In brief, the CTMC layer computes the log-likelihood *L*_*mi*_ of each alignment column *i* ∈ {1, *…, 𝓁*} simultaneously in several evolutionary models *m* ∈ {1, *…, M*} (see Figure 1). We assume the alphabet has *s* characters that are numbered 1, *…, s*, in our application the alphabet is the set of *s* = 64 codons **{** aaa, aac, *…*, ttt**}**, i.e. each codon is a single character. However our program ClaMSA is not specific for codon evolution. Neither does the model explicitly use a genetic code.

**Figure 1.**
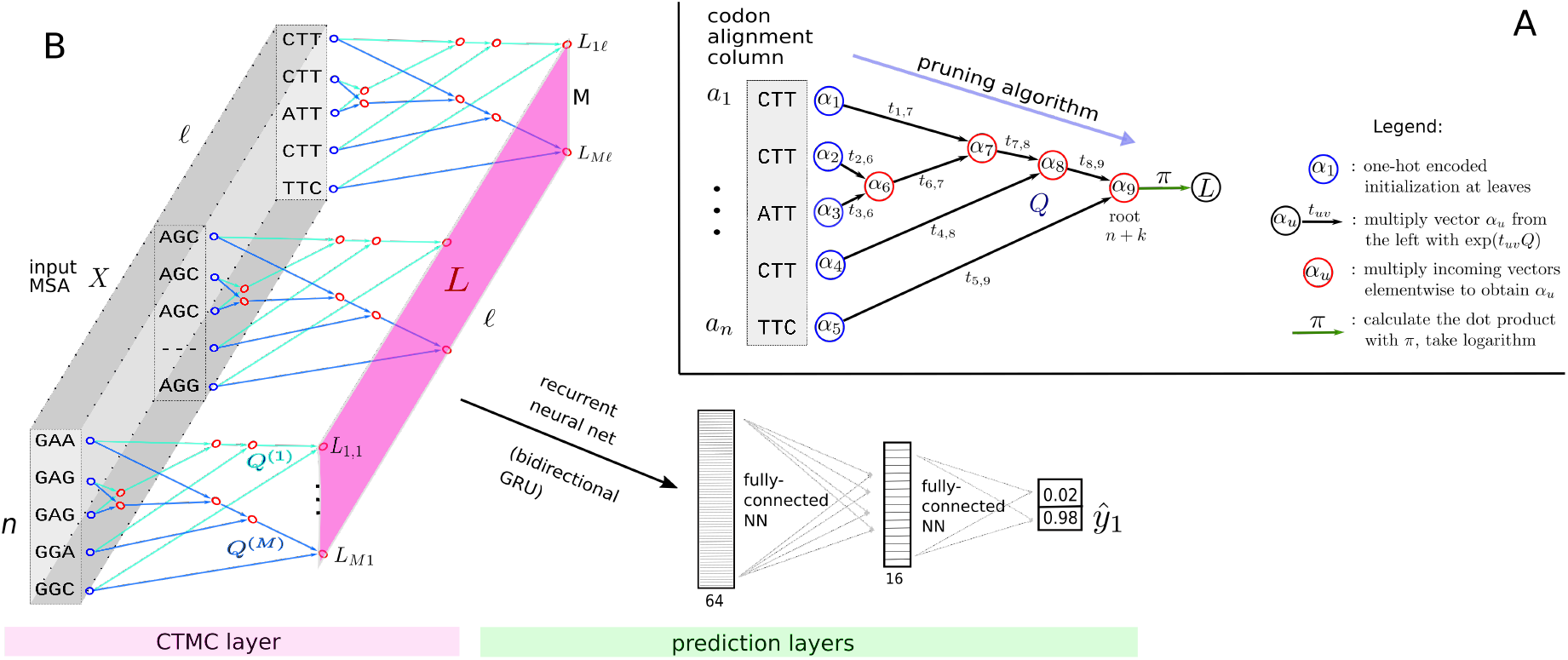
CTMC layer and overall model. **A**: A dynamic programming algorithm computes in a leaf- to-root traversal of the tree, the log-likelihood *L*, i.e. the logarithm of the probability of an alignment column (blue leaves), marginalizing over all unknown characters at the ancestral nodes (red). **B:** Log-likelihood *L*_*mi*_ is computed for each of *M* phylogenetic models specified by rate matrices *Q*^(1)^, *…, Q*^(*M*)^ and for each alignment column 1, *…, 𝓁*. The remaining layers (Gated Recurrent Unit, Dense) estimate the probability *y*_1_ that the input alignment is coding.

We choose a standard parametrization of GTR *s* ×*s* rate matrices, except that we insert another first dimension to account for *M* different evolutionary models. Different evolutionary models *m* = 1, 2, 3, *…* could e.g. represent protein-coding, non-coding, conserved non-coding and other subsets of aligned sequences. However, *m* can be interpreted as a *latent* variable and the MSAs are not explicitly assigned to a particular evolutionary model. Let the rate matrices *Q* be defined as

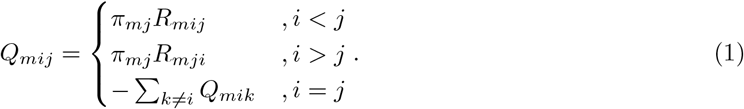

Here, *m* indexes an evolutionary model, π ∈ [0, 1]^*M* × *s*^ are the *stationary probabilities* of the *M* GTR models (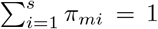 for all *m*) and *R* ∈ ℝ_+_ ^*M* × *s* × *s*^ holds the rate factors: For each model *m* the matrix *R*_*m*,:,:_ is upper triangular and holds *s*(*s* − 1)*/*2 positive rate factors. Here, we use a colon notation analogous to Python code `R[m,:,:]` of Numpy, TensorFlow or PyTorch. We also use the notation *Q*^(*m*)^ = (*Q*_*mij*_)_*ij*_ for the *m*-th rate matrix.

The *input* of the CTCM layer are multiple sequence alignments, specified as a 3-tensor *X* ∈ {0, 1}.^*𝓁 × n × s*^ in one-hot encoding, i.e. *X*_*iua*_ = 1 if and only if sequence *u* has character *a* at site *i*. Gaps and completely unaligned sequences are treated separately as missing data. Further, the layer requires a tree *T* with *times* or branch lengths *t* that relates the aligned input sequences. In our experiments we used the same species tree for all MSAs in a given clade. The tree can be unrooted but we assume that it is rooted. The choice of the root is arbitrary and without effect on the output as all Markov chains are by construction time-reversible. We do not require that each ancestral node has exactly two children. Let *T* = ({1, *…, n* + *k*}, *E*) be the directed tree with *n* leaves 1, *…, n*, with *k ancestral nodes n* + 1, *…, n* + *k* (non-leaves) and with edges *E* that are directed towards the *root* node *n* + *k*. For an example, see Figure 1 A).

The output of the CTMC layer is an *M* × *𝓁* matrix *L* = (*L*_*mi*_), where entry *L*_*mi*_ is the log-likelihood of the *i*-th alignment column (‘site’) under the *m*-th evolutionary model:

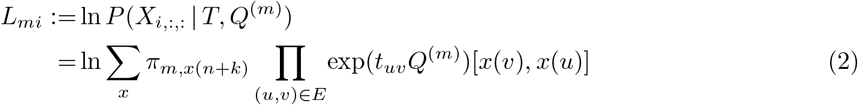

Here, *x* : {1..*n* + *k*} *→* {1..*s*} ranges over all assignments of nodes of tree *T* to characters that agree on the leaf set {1..*n*} with the observed characters (*X*_*iua*_ = 1 ⇒ *x* (*u*) = *a*), *t*_*uv*_ is the length of the branch from node *u* to node *v*, exp is the matrix exponential and the brackets [,] in (2) are a notation to index the exponentiated matrix in row *x*(*v*) and column *x*(*u*).

#### 3.1.1 Pruning Algorithm

It is well-established [Felsenstein, 1973] that the likelihood of the leaf data, implicitly given by equation (2), can be computed for constant alphabet size *s* in time *O*((*n* + *k*)*𝓁*) with the pruning algorithm, a dynamic programming algorithm that computes conditional likelihoods of subtrees in a leaf-to-root order. Algorithm 1 is essentially the execution of this pruning algorithm in parallel for all models *m* and all sites *i. α* is the array of dynamic programming variables: *α*_*m,i,v,a*_ is the conditional likelihood under the *m*-th phylogenetic model of the leaf assignments at site *i* in the subtree of *T* rooted at node *v*, given that node *v* is assigned character *a*. The algorithm makes use of vectorization and *shape broadcasting*. Line 5: exp is the matrix exponential, here applied to all square matrices *t*_*u,v*_*Q*^(*m*)^ for all *m* in parallel and the resulting matrices are stacked, so that the red array (tensor) is again of shape *M* ×*s* × *s*. The multiplication of the red and blue factors are *M* × *𝓁* multiplications of a *s* ×*s* matrix with a size *s* vector (shape broadcasting). Line 7: The multiplication is a dot product simultaneously carried out for *M* × *𝓁* pairs of size *s* vectors and the logarithm is applied elementwise.

##### Algorithm 1 CTMC(*X*; *π R*; *T*).

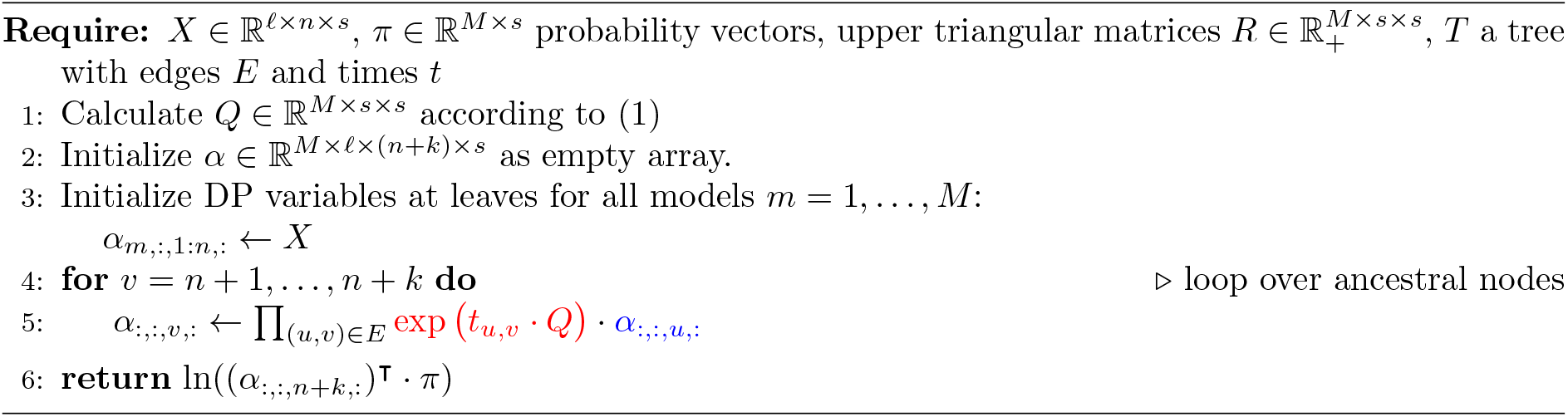

## 3.1.2 Learning CTMC

Let Θ := (*π, R*) be the parameters of the CTMC model that need to be learned. Because the CTMC layer is designed to be used as one of several layers that can be learned end-to-end using stochastic gradient descent, we need to be able to compute the gradient

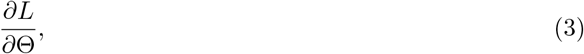

where *L* = (*L*_*mi*_) = CTMC(*X*; *π, R, T*) is the output of the layer. This was achieved using the automatic differentiation capabilities of TensorFlow [Abadi et al., 2016]. In particular the gradient of the matrix exponential *exp*(*A*) = *I* + *A* + *A*^2^*/*2 + *A*^3^*/*3! + … is required. Like in Matlab and R, in TensorFlow the matrix exponential is approximated using the Padé approximation [Higham, 2005]. This approach requires as elementary functions only the multiplication and addition of matrices as well as computing an inverse and a power of a matrix. These functions are all automatically differentiated by the TensorFlow backend.

For the default choice of *M* = 8 phylogenetic models and the codon alphabet *s* = 64 the CTMC layer has 16632 = 8 × (63 + 64 *·* 63*/*2) parameters to store eight rate matrices implicitly through stationary distributions *π*_*m*,:_ and triangular rate factor matrices *R*_*m*,:,:_ each.

### 3.2 Prediction Layers and Loss

The output *L* of the CTMC layer has dimensions of sizes *M* and *𝓁*, of which the alignment length *𝓁* varies with the input. In order to produce a fixed-dimension output, e.g. the probabilities of two classes, regardless of the input length, we add layers that aggregate along the sequence dimension and produce a prediction. We will refer to the layers after the CTMC layer the *prediction layers* (see Figure 1).

### Sequence Model: Recurrent Neural Network

By default ClaMSA uses a *sequence model* and fully-connected neural network for binary classification. To describe the architecture we will use high-level machine-learning notation like `L = CTMC(X),` which means that *L* is the output that the CTMC layer produces on input *X*. The default version of ClaMSA uses a Gated Recurrent Unit (GRU) as sequence layer [Cho et al., 2014]. As a special kind of recurrent neural network, a GRU is, in principle, capable of modeling autocorrelation that MSAs can show, e.g. due to local secondary or tertiary structure. Finally, two “Dense” layers, which are fully-connected neural networks (sometimes called multilayer perceptrons) produce the prediction:

~~~
rnn = Bidirectional(GRU(32))(L)
D = Dense(16, activation = “sigmoid”)(rnn)
y = Dense(2, activation = “softmax”)(D)
~~~

These prediction layers have 9138 = 8064 (bidirectional GRU for *M* = 8) + 1040 (first Dense) + 34 (second Dense) parameters. Here, half of the parameters of the output (last) layer are redundant and could be saved using the logistic sigmoid function as activation for ŷ. However, this form allows ClaMSA to be used for more than two output classes. The suitability for multiclass classification is an advantage over methods that use a likelihood ratio (PhyloCSF) or a threshold on ŵ= *dN/dS* (PAML). The training criterion is to minimize the cross-entropy error (CEE) as loss function

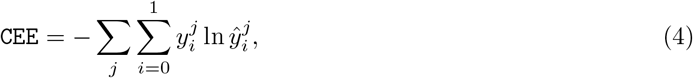

where *ŷ*^*j*^ and *y*^*j*^ ∈ {0, 1}^2^ are the vector of the two predicted probabilities and the corresponding one-hot encoded *training label* for the *j*-th training example, respectively. All rate matrices as well as the parameters of the prediction layers are trained simultaneously with stochastic gradient descent to mimize the loss on labeled training data.

### Logistic Regression

In order to be able to attribute performance gains to the CTMC layer rather than the prediction layers, we also implemented a very simple alternative that we call the *LogReg* variant of the prediction layers. A simple averaging of the *M* log-likelihoods along the sequence dimension (of length *𝓁*) produces a sequence model output of fixed size *M*.

~~~
means = mean(L, axis = 1)
y = Dense(2, activation = “softmax”)(means)
~~~

These prediction layers have only 2(*M* +1) = 18 parameters (*M* +1 = 9 free parameters) and compute after a parameter transformation the following prediction function:

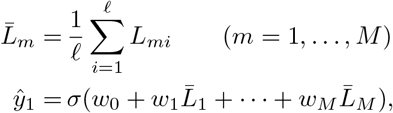

where σ (*z*) = 1*/*(1 + exp(— *z*)) is the logistic sigmoid function. Setting *ŷ*_0_ := 1— *ŷ*_1_ we again use CEE as optimization criterion

### Maximum-Likelihood

In principle, the CTMC model also allows to compute a maximum likelihood estimate of a rate matrix under a model of independent alignment columns. This can be achieved when setting *M* = 1 and using 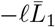 as loss funtion. We confirmed on simulated alignments that indeed a ML-estimate of *Q*^(1)^ approximated well the rate matrix used in the simulation (data not shown), but have not used this functionality in the work described here.

#### 3.3 Data Sets

For training and testing ClaMSA we generated multiple sequence alignments of potential coding exons (‘exon candidate MSAs’) in a clade of vertebrate species, a clade of fly species and a clade of yeast species as summarized in Table 1. The first two clades were previously often used to benchmark comparative genomics methods. The yeast clade was chosen because it is the only non-metazoan clade for which PhyloCSF provides parameters. In each clade, the set of labeled MSA examples was randomly partitioned to achieve a defined size of the test and validation set, the remaining examples constitute the training set.

**Table 1:**
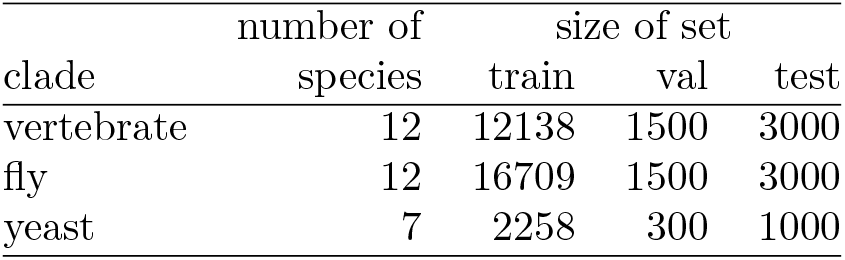
Data sets of aligned candidate exons.

| clade | number of | size of set |  |  |
| --- | --- | --- | --- | --- |
|  | species | train | val | test |
| vertebrate | 12 | 12138 | 1500 | 3000 |
| fly | 12 | 16709 | 1500 | 3000 |
| yeast | 7 | 2258 | 300 | 1000 |

As basis for finding alignments of candidate coding regions we used multiple genome alignments. For the vertebrate clade we downloaded a MULTIZ multiple genome alignment from the UCSC Genome Browser [Lee et al., 2020], for fly and yeast we constructed one ourselves using CACTUS [Paten et al., 2011]. An exon candidate MSA is an aligned tuple of potential exons in the same reading frame that does not contain a stop codon in any of the considered rows. Each row could therefore be a complete coding exon. Exon candidate MSAs were compiled as described in the supplementary data and labeled as coding (positive, *y*_1_ = 1) or noncoding (negative, *y*_1_ = 0) using a reference genome annotation. The negative examples were filtered not to overlap a reference coding region in the same frame and on the same strand, as such a case could arguably be counted as partially correct. For practical reasons and as the vertebrate and fly data sets are strongly imbalanced towards the negatives labels, we chose a ratio of 1:2 between positive and negative examples.

The negative exon candidate alignments are on average shorter than the positives. In order to remove the influence of the length, we subsampled the negative examples by length, so that for each species the positives and negatives have very similar length distributions (see Supplementary Figure 1). The average number of exon candidates (= rows, species) in an candidate exon MSA are 4.5 (vertebrate), 4.1 (fly) and 3.0 (yeast). The average length in nucleotides are 158, 367 and 688, respectively.

#### Tree construction

A random selection of positive exon candidate MSAs was converted with `ClaMSA` convert to NEXUS format and subsequently a tree constructed with MrBayes [Ronquist et al., 2012] for each clade. The three trees are show in Supplementary Figure 3, as well as the command lines. They were used as input to the training and prediction procedures.

#### Codon alignments

As a preprocessing, nucleotide alignments with an assumed reading frame for all sequences of the MSA are converted to codon alignments as shown in Supplementary Figure 2. Thereby, two codons are aligned with each other in the output codon alignment if and only if all three corresponding bases were aligned with each other in the input nucleotide alignment.

## 4 Results

### 4.1 Benchmarking of Tools

We compared ClaMSA against PhyloCSF [Lin et al., 2011] and the program codeml from the PAML package [Yang et al., 1997]. PhyloCSF and `ClaMSA` were input identical FASTA formatted multiple sequence DNA alignments that started with complete codons of a candidate coding region. The frame or strand needed not be inferred by the programs. `Codeml` requires codon alignments as input. Therefore, for its input the nucleotide alignments were first converted to codon alignments with `clamsa convert.` The command lines for running the programs are stated in the supplementary data.

### Evaluation as a binary decision

Each tool outputs a number: ClaMSA a probability, PhyloCSF a log-likelihood ratio and `codeml` an estimation for *ω* (dN/dS rate ratio) (see Suppl. Figure 4). In each case the number can be interpreted as a binary decision on whether a given input candidate exon alignment is coding in the specified frame and on the specified strand. For ClaMSA the prediction is positive if and only if (iff) the output probability of “coding” is larger than 50%, for PhyloCSF iff the log-likelihood ratio is larger than 0, for `codeml` iff the estimated *ω* is less than 1, i.e. if the sequences are estimated to have been as a whole under purifying (negative) selection.

Table 2 shows the results on the test sets summarized in Table 1. Method “ClaMSA” was trained on a training set of alignments from fly, vertebrate and yeast sequences, mixed at equal parts. “ClaMSA{fly, vert, yeast}” refer to a version that was trained on a training set from the respective clade. Column “errors” shows the total numbers of misclassification (false positives or false negatives). In all three clades ClaMSA made the fewest errors, PhyloCSF made about 7, 2.5 and 7 times as many errors on vertebrates, flies and yeasts, respectively, `Codeml` even more (Table 2, Suppl. Tab. 1). On each clade we compared the default ClaMSA which had been trained on a mixture of alignments from the three clades with a *native* version that was trained, like PhyloCSF, on the same clade the test is performed on. For the vertebrate and fly clades this version of ClaMSA was somewhat or considerably better than the universal version that was otherwise used for experiments on all clades.

**Table 2:**
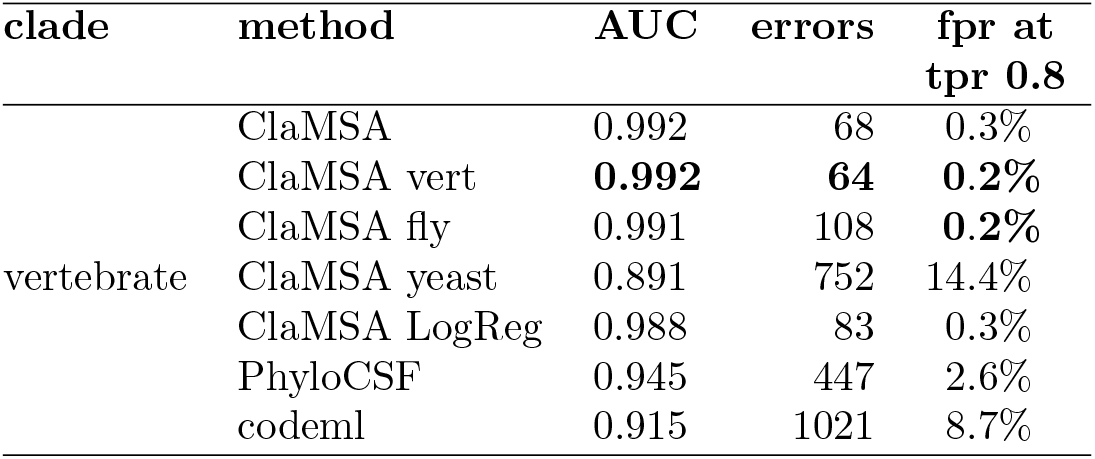
Accuracy Comparison. Best performances are in bold face.

### ROC analysis

Above thresholds (0.5, 0, and 1 for ClaMSA, PhyloCSF and `codeml`, respectively) need not be reasonable, e.g. when in a genome-wide scan a large number of false positives is not tolerable or if – contrary – a very sensitive search is required. We therefore also compared the receiver operating characteristic (ROC) curves. Table 2 shows the area under curve (AUC) values and in the last column the false positive rate (false positives/actual negatives) at the respective choice of the threshold for which the true positive rate (sensitivity, true positives/actual positives) is just above 80%. The difference between the AUC and the optimal value of 1, is reduced by ClaMSA in comparison to PhyloCSF from 5.5% to 0.8% on vertebrates, from 9.5% to 2.1% for flies and from 4.4% to 2.1%. Apparently, the accuracy advantage of ClaMSA over PhyloCSF is smaller for yeast than for the other two clades. A possible explanation is that the overall number of exons is small in yeast as most genes are intronless. In that sense yeast constitutes an atypical example for the commonly studied eukaryotic genomes because most CDS are complete genes and can therefore more easily be identified with non-evolutionary methods. We compiled only a total of 4590 positive examples of aligned coding exons in yeast. It can be expected that the overlap of the training set of PhyloCSF with our random test sets makes up a larger fraction of the overall training set of PhyloCSF that is not available to us. We included results on yeast in order to be able to compare to PhyloCSF on a non-animal clade.

### Influence of alignment attributes

Counterintuitively, all tools – and most strikingly PhyloCSF – appear to loose accuracy when the MSAs are particularly long (see Supplementary Figure 5). This is mainly due to false positive classifications of long alignments of few (mostly two) sequences. It appears that that section of the data could be classified with fewer errors using a simple additional filter on MSA width and length. We chose not to use the width or length of the alignment as a criterion as this would likely degrade the generalizability across clades. In addition, we expect that arguably most search tasks for novel coding regions would filter out candidate exons that are present in only very few species, anyways.

### 4.2 Generalizability and Cross-Clade Training

Both ClaMSA and PhyloCSF train 64 × 64 codon rate matrices with thousands of parameters, which raises the question of generalizability: Can the accuracy on users’ sequence data ‘unrelated’ to the training set be expected to match the average accuracies reported here? This question will be discussed on several levels.

Firstly, an overlap between our test sets and the training sets could account for a lower performance than reported here. For ClaMSA we have naturally excluded any overlap. However, as PhyloCSF cannot be trained by users, we could not control the fraction of test genes that were also used in its training. This could in particular be an issue for the yeasts as their genomes have in comparison very small numbers of exons.

Secondly, homologs of any degree of sequence similarity can exist within the genome of a species and between species. Data that is guaranteed to be ‘unrelated’ to the training data cannot be found. More important is, however, that the parameter sets perform well when used to classify alignments from a broad set of species, e.g. because intrinsic properties of the genetic code and amino acid similarities were learned, rather than the frequencies of individual transitions events in the training set.

We therefore tested, how well the parameters of ClaMSA generalize between clades. Table 2 shows these cross-clade applications. For example, when ClaMSA was trained on fly alignments only (ClaMSA fly) its performance on vertebrates is with an AUC of 99.1% almost as good as the version that was natively trained (ClaMSA vert, AUC 99.2%). Vice versa, when ClaMSA was trained on vertebrate MSAs only it performed somewhat worse (AUC 95%) than ClaMSA that was trained on flies (AUC 97.9%), but still considerably better than PhyloCSF (AUC 90.5%) (Supp. Table 1). Note for these comparisons that, we always used the PhyloCSF version that was trained for the clade we tested its performance on. Thusly, the cross-species performance of ClaMSA is in these cases better than the within-clade performance of PhyloCSF. However, this is not the case for cross-clade training between an animal and yeast. The vertebrate and fly version perform much worse on yeasts. However, the default ClaMSA version which was trained on all three clades performs even somewhat better than the native version (Supp. Table 1).

### 4.3 Breakdown of Accuracy Gains

This manuscript contains several new ideas for approaches, that more traditional approaches for estimating codon rate matrices such as PAML and PhyloCSF do not have; in particular, a variable number of empirical rate matrices, prediction layers (sequence model) that have their own parameters, end-to-end learning and discriminative rather than generative learning of the matrices. In order to assess the individual contribution of some of the main ideas to the overall accuracy, we have compared ClaMSA against versions of itself, where some component has been replaced or parameters have been changed.

The number of rate matrices *M* has only a small influence on the accuracy, provided it is at least 3 (Supplementary Figure 6), suggesting that most of the gain is not due to the larger number of parameters of the model.

We replaced the gated recurrent unit (GRU) as sequence model against the very simple logistic regression model introduced in section 3.2, called ‘ClaMSA LogReg’ in table 2. It is otherwise trained like the default version of ClaMSA. Omitting the recurrent neural network reduces the AUC on vertebrates and fllies only by 0.4 and 1 percent points, respectively. The reduction on yeasts is more pronounced (94.3% versus 97.9% AUC). A possible explanation is that yeast exons are usually complete genes and therefore a bidirectional recurrent neural network could benefit more from learning positional inhomogeneity than on clades where introns can rather arbitrarily interrupt coding sequences.

Further, we compared the GRU against a simple RNN as well as a long short-term memory (LSTM) model. For each sequence model class we varied the number of units. Both alternative sequence model classes performed slightly worse than the GRU (data not shown).

### 4.4 Application Example

Figure 3 shows an example of a candidate novel human coding region that was identified with ClaMSA. The aligned genome regions in cow and tenrec also have no gene annotated. However, the presumably homolog regions in mouse and rat have annotations in GENCODE and RefSeq. The homolog of the region depicted in Figure 3 is a large initial CDS of the mouse gene Zpld2 (Supplementary Figure 7).

**Figure 2.**
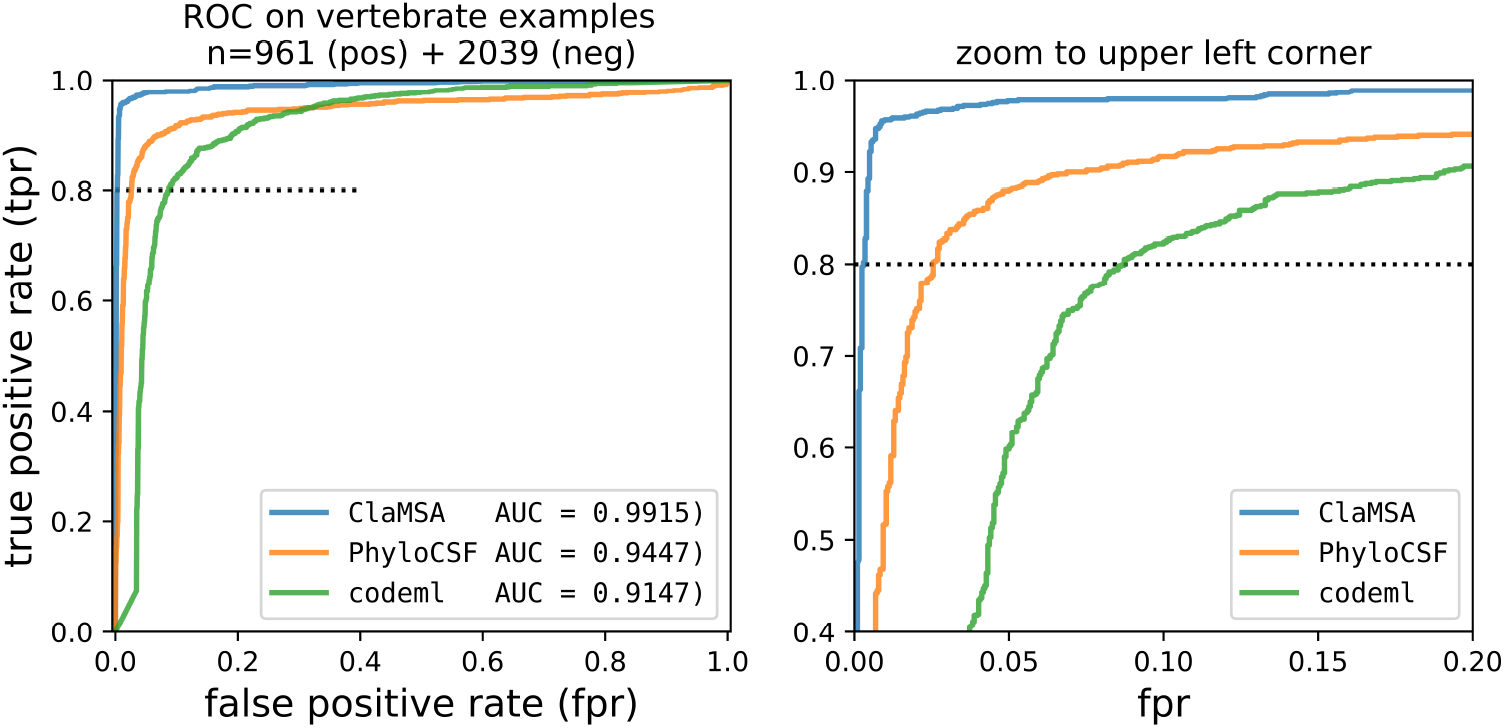
Accuracy on vertebrates.

**Figure 3.**
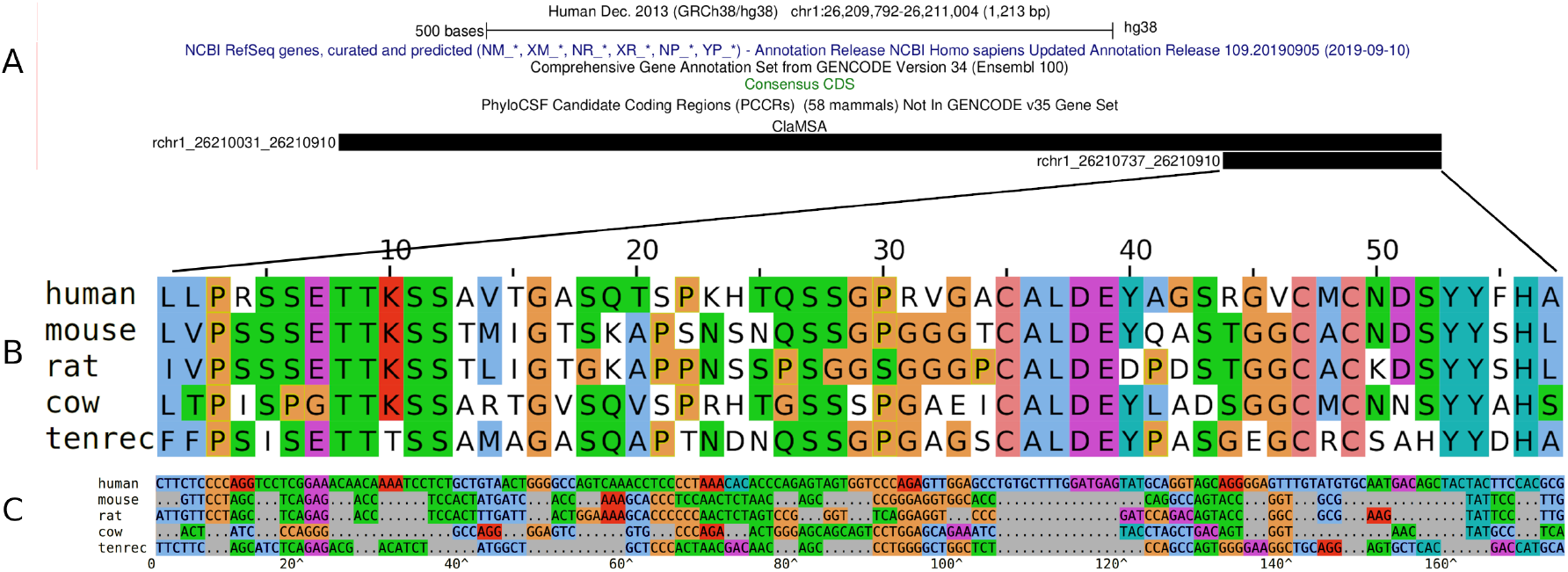
Discovery of a presumably novel human coding region with ClaMSA. **A**: A region in the human genome that contains two candidate exon MSAs (black bars) that were highly confidently (94.9% and 93.3% probability) classified by ClaMSA as coding. The regions are bordered on both sides by conserved splice site patterns, suggesting the larger ClaMSA region is part of a novel human spliced gene. The UCSC genome browser [Lee et al., 2020] contained in March 2021 neither an annotation from RefSeq, GENCODE or CCDS, nor were there aligned transcript sequences, retroposed genes or repeats (Supplementary Figure 7). There is also no PhyloCSF candidate region [Mudge et al., 2019]. **B**: Amino acid alignment. The alignment in the regions of the smaller ClaMSA region is gapless. **C**: Codon alignment. Codons that are identical to human are shown as “…” and in gray. Codons differing from the human codon are shown explicitly and are color-coded by groups of similar amino acids. codeml estimates a rate ratio of nonsynonymous versus synonymous mutations of *ω* = dN/dS = 0.61, suggesting the region has been under purifying selection.

## 5 Discussion

We applied a modern machine learning approach to estimate the parameters of a classical statistical model that has been used in phylogenetics for many decades. With it, the state-of-the-art performance in classifying multiple alignments phylogenetically can be pushed significantly. We demonstrated that the binary classification of codon multiple sequence alignments into coding and non-coding can be much improved. The previously published method PhyloCSF was based on a ratio of likelihoods of two generatively and independently trained rate matrices. Our new tool ClaMSA, however, trained all rate matrices simultaneously discriminatively and end-to-end together with a prediction model. On our test set of 12-way vertebrate alignments, ClaMSA reduced the false positive rate of 2.6% of PhyloCSF to 0.2% when classifying 12-way vertebrate alignments at sensitivity levels of 80%, *even when its rate matrices were trained on flies*.

## 6 Conclusion

ClaMSA does not necessarily require pre-trained parameters for each specific clade. Overall, we believe that most users may prefer to use the default parameters that performed well in all our experiments over retraining parameters for a specific clade of interest. However, ClaMSA provides the functionality for users to train it given labeled alignments and a tree. Possible direct applications of ClaMSA are the classification of candidate ORFs as shown here, based on this a genome-wide scan for novel coding regions, or a quantitative genome browser track with per base coding probabilities.

Our discriminatively trained rate matrices can be used by other tools as well. Tools that use a rate matrix and the pruning algorithm to compute the log-likelihood of a codon site can replace their own matrix by the ones we trained. We have published for this purpose a set of rate matrices in text format that is included in the ClaMSA distribution. In particular, we plan to use these ClaMSA-trained codon rate matrices for comparative gene prediction with AUGUSTUS [König et al., 2016]. Other potential applications are the evolutionary discrimination of amino acid or DNA alignments.

Even of higher meaning could be the general CTMC layer that allows to compute gradients of the tree-likelihood under the almost universally used continuous-time Markov chain model. The differentiation can be done with respect to the rate matrix as well as simultaneously to the branch lengths (not used here). In contrast, common practice in phylogenetic programs is the numerical differentiation with respect to each rate matrix parameter individually (RAxML, Stamatakis [2014]) or to simultaneously make small random changes to all rates (MrBayes, Ronquist et al. [2012]).

## Acknowledgements

This work was supported by the Swiss National Science Foundation [Gesuch Nr. 407540 167331 to M.S.].

We thank Lizzy Gerischer for making initial comparisons between PhyloCSF and an *ω*-based classification method.

