## Supplementary Materials for "End-to-end Learning of Evolutionary Models to Find Coding Regions in Genome Alignments"

Darvin Mertsch[1] and Mario Stanke[1,2]

[1]Institute for Mathematics and Computer Science, Universität Greifswald, Walther-Rathenau-Str. 47, 17489, Greifswald, Germany
[2]Center for Functional Genomics of Microbes, Universität Greifswald, 17489 Greifswald, Germany

### 1 Supplementary Methods

#### 1.1 General Time-Reversible CTMC on a Tree

Let $\Sigma = \{1, \ldots, s\}$ be a finite state space and let $T$ be a tree with $n$ leaves $1, \ldots v$, $k$ inner nodes $n+1, \ldots, n+k = \mathrm{root}(T)$, the edges being directed towards the root and with branch lengths $t_{uv} > 0$ for any edge from node $u$ to any node $v$. We call a family of $\Sigma$-valued random variables $a_1, \ldots, a_{n+k}$ a *general time-reversible (GTR) model on $T$* if and only if there exists a stationary, homogeneous and time-reversible continuous-time Markov chain $X$ on $\Sigma$ such that

- $\mathbb{P}[a_u|a_v] = \mathbb{P}[X_{t_{uv}} = a_u|X_0 = a_v]$ for all edges $(u, v)$ of length $t_{uv}$,
- $\mathbb{P}[a_{n+k}] = \pi$, where $\pi$ is the stationary probability vector of $X$.

In the classical model of Tavaré (1986) one would take $\Sigma = \{1, 2, 3, 4\}$ corresponding to $\{\mathtt{a}, \mathtt{c}, \mathtt{g}, \mathtt{t}\}$ in order to model evolution in the state space of nucleotides. For our purposes we choose $\Sigma = \{1, \ldots, 64\}$ to model evolution in the state space of codons $\{\mathtt{aaa}, \mathtt{aac}, \ldots, \mathtt{ttt}\}$. The alignment columns of an MSA are interpreted as (one-hot encoded) leaf configurations for the random variables $a_1, \ldots, a_n$. Given such a GTR model on $T$ we want to calculate the likelihood $L = \mathbb{P}[a_1, \ldots, a_n]$ of any leaf configuration. In brief, our CTMC layer calculates these likelihoods in a trainable way (with respect to the Markov chain). More precisely, our CTCM layer is configurable to simultaneously store $M$ independent GTR models. The Markov chains $X^{(m)}$ of these GTR models are represented by a parameterization of their respective rate matrices $Q^{(m)}$ by a probability vector $\pi^{(m)}$ and a strict upper triangle matrix $R^{(m)}$ with positive entries.

#### 1.2 Alignment Data Sets

**Clades, reference annotations and genome alignments.** We generated labeled alignments from original input. In each clade we used a reference annotation of one species. For the *vertebrate* clade (human, rhesus, mouse, rat, rabbit, cow, dog, elephant, tenrec, armadillo, opossum, chicken) we used the RefSeq annotation of coding exons (CDS) of the human genome. For the *fly* clade (*Drosophila melanogaster*, *D. simulans*, *D. sechellia*, *D. erecta*, *D. yakuba*, *D. ananassae*, *D. virilis*, *D. mojavensis*, *D. grimshawi*,*D. persimilis*, *D. pseudoobscura*, *D. willistoni*) we used the FlyBase annotation of *D. melanogaster.* For the yeast clade (*Saccharomyces cerevisiae*, *S. paradoxus*, *S. mikatae*, *S. kudriavzevii*, *S. bayanus*, *S. castelli*, *S. kluyveri*) we used the RefSeq annotation of *S. cerevisiae.* For the vertebrate clade we downloaded a MULTIZ multiple genome alignment from the UCSC Genome Browser (Tyner et al., 2016), for fly and yeast we constructed one ourselves using CACTUS (Paten et al., 2011).

**MSA construction.** Exon candidate MSAs were generated with comparative AUGUSTUS (König et al., 2016). AUGUSTUS-CGP first identifies potential coding exons ('exon candidates') in each of the aligned genomes. An exon candidate is a tuple (`species`, `seqname`, `type`, $a$, $b$, $\epsilon$, $f$), where `type` specifies the signals expected at the boundaries, e.g. internal exons are bordered by a donor and an acceptor splice site, $a \leq b$ are the interval boundaries, $\epsilon \in \{-1, +1\}$ is the strand and $f \in \{0, 1, 2\}$ is the reading-frame. AUGUSTUS-CGP identifies all exon candidates without an in-frame stop codon whose boundaries score a minimum threshold in their respective model (e.g. splice site model) or that were sampled by the AUGUSTUS gene prediction model or that align at both boundaries with an exon candidate from another genome ('liftover'). When multiple exon candidates are aligned at both boundaries and agree on type, frame and strand, they are summarized as an *exon candidate MSA*, thereby reducing the genome alignment to the exon candidates. The command line for producing these exon candidate MSAs are documented in the file `EXONCAND-MSAS-CGP.md` of the AUGUSTUS package. An exon candidate MSA is considered *positive* if it contains an exon candidate that *exactly matches* a coding exon from the reference annotation. From the remaining exon candidate MSAs those were removed that overlap an exon from the reference annotation on the same strand and in the same frame. The *negative* exon candidate MSAs therefore do not overlap an exon from the reference annotation, or they overlap but are in a different reading frame or strand.

**Downsampling and random split.** The negative exon candidate alignments are on average shorter than the positives. In order to exclude their length as a criterion we subsampled the negative examples by length, so that for each species the positives and negatives have very similar length distributions. Further, for practical reasons we chose a probability ratio of 1:2 when sampling positive and negative examples, as the vertebrate and fly data sets are strongly imbalanced towards the negatives labels. Subsequently, for each data set the examples are randomly permuted under the uniform distribution and the examples were partitioned into a training, validation and test split as shown in Table 1 of the main document.

#### 1.3 Subsampling to Harmonize Length Distributions

Initially, the negative examples had a length distribution that significantly deviates from that of the positive examples (negatives have much smaller mean). For that reason the negative examples were subsampled based on the length only so that the training, validation and test sets had very similar length distributions.

#### 1.4 Codon Alignments

As a preprocessing, nucleotide alignments with an assumed reading frame for all sequences of the MSA are converted to codon alignments as shown in Figure 2. Thereby, two codons are aligned with each other in the output codon alignment if and only if all three corresponding bases were aligned with each other in the input nucleotide alignment.

#### 1.5 Tree Construction

The trees shown in Supplemenary Figure 3 were all constructed the same way. We consider the uniformity of construction a prerequisite to the transfer of parameters across clades. The basis for tree construction were exon candidate MSAs constructed with comparative AUGUSTUS as described in the file `EXONCAND-MSAS-CGP.md` of the AUGUSTUS distribution.

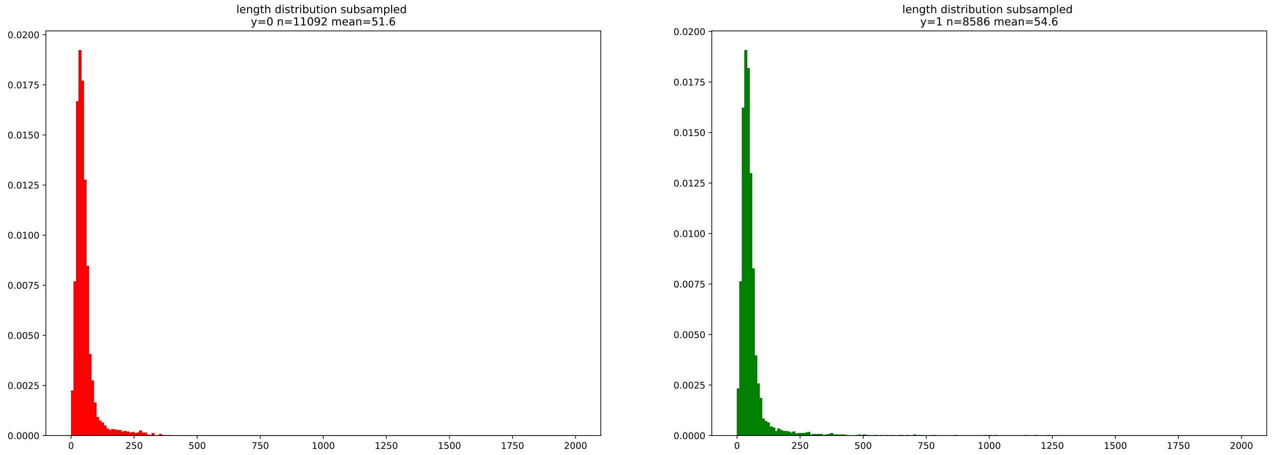

**Supplementary Figure 1:** Subsampling negative examples. Non-coding MSAs (red, left) with overrepresented lengths were downsampled so the length is by itself not a discriminating feature. Here, the numbers of codons from the vertebrate clade are shown.

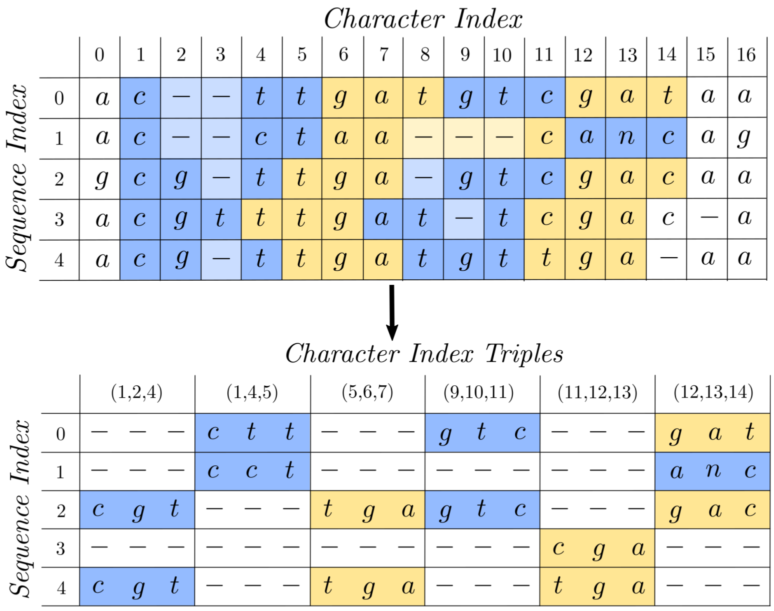

**Supplementary Figure 2:** Example how an input *nucleotide* alignment on the plus strand with a global frame of $f = 1$ (top) is preprocessed and converted to a *codon* alignment (bottom). Codons are color-marked.

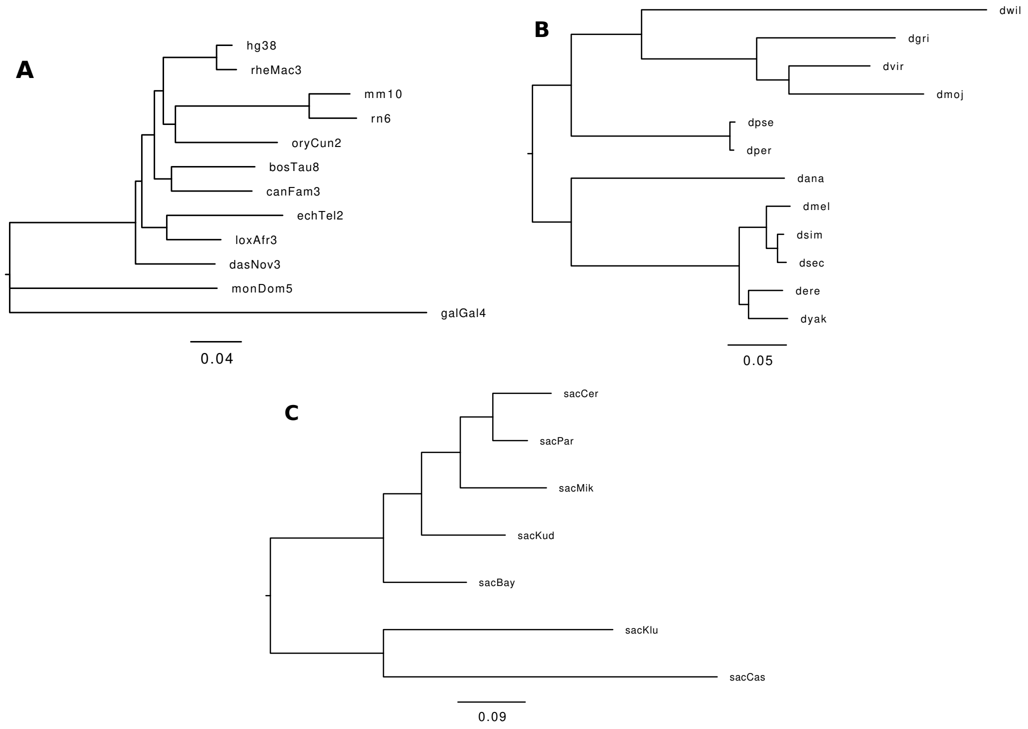

**Supplementary Figure 3: Phylogenetic trees of the used clades** A: vertebrate, B: fly: C: yeast. The trees were constructed with MrBayes (Ronquist et al. (2012)) from nucleotide alignments of coding sequences and are scaled to expected number of nucleotide mutations per site.

The following command lines assume that this output has been stored in gzip compressed files `aug*.out.gz`. They are given for the example of yeast.

```
# convert nucleotide alignments to codon alignments in NEXUS format
clamsa.py convert  augustus aug*.out.gz --use_codons --margin_width 0 \
  --write_nexus yeast.nex --clades tree-phylocsf.nh --nexus_sample_size 500

# run MrBayes (here in parallel) with a codon model
# lset nst=6 Nucmodel=Codon omegavar=M3 rates=gamma;
mpirun -use-hwthread-cpus -np 4 mb yeast.nex
# took about 15 hours
```

#### 1.6 Running Classifiers

**Running ClaMSA** Predictions were done in parallel for different training configurations (e.g. choices of $M$, training clades, prediction models)

```
for clade in vertebrate fly yeast; do
    clamsa.py predict fasta \
        $clade.test.msa.lst \
        --clades fly.nwk vertebrate.nwk yeast.nwk \
        --use_codons --batch_size 20 \
        --log_basedir ...  --saved_weights_basedir ... \
        --model_ids '{
            "fly_vert_yeast-rnn8":  "2020.12.24--21.23.47",
            "fly_vert_yeast-mean8":  "2020.12.25--03.23.32",
            "fly-rnn8":  "2020.12.24--02.20.16",
            "fly-mean8":  "2020.12.24--12.34.50",
            "vert-rnn8":  "2020.12.24--05.26.08",
```

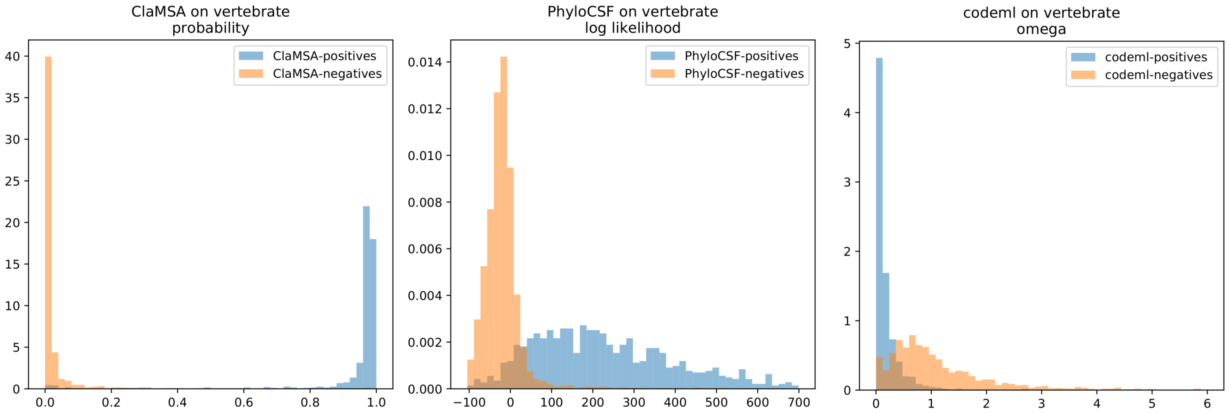

**Supplementary Figure 4:** Distribution of vertebrate outputs. The output is either a probability (ClaMSA, left), a log-likelihood ratio (PhyloCSF, middle) or an estimate of dN/dS (codeml, right). The outputs of the positives (blue) and negatives (orange) are fairly well separated with ClaMSA. For the other two tools the two distributions have a clearly visible overlapping region, such that any threshold will lead to a number of false (positive or negative) classifications.

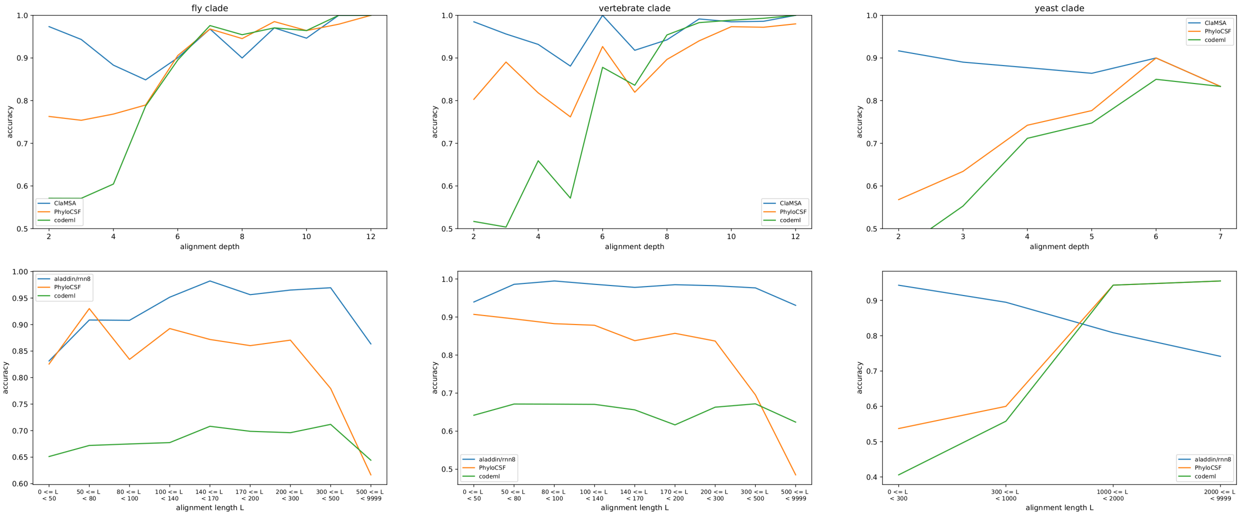

**Supplementary Figure 5:** Accuracy as a function of alignment depth and length. The depth is the number of aligned sequences. The length is the number of alignment columns.

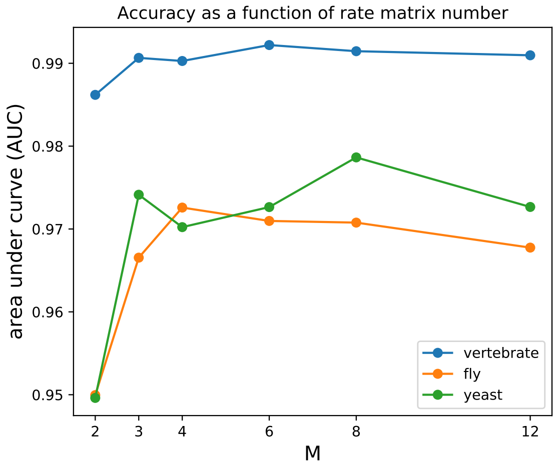

**Supplementary Figure 6:** Choice of number of evolutionary models $M$.

```
      "vert-mean8":  "2020.12.24--11.34.50",
    ...
  }' \
--out_csv preds_${clade}_test.csv
```

**Running PhyloCSF:**

```
PhyloCSF --removeRefGaps -f 1 100vertebrates msa.fa
PhyloCSF --removeRefGaps -f 1 12flies msa.fa
PhyloCSF --removeRefGaps -f 1 7yeast msa.fa
```

`codeml` **configuration:**

```
  seqfile = msa.fa * sequence data filename
 treefile = tree.nwk * tree structure file name
  runmode = 0  * user tree
  seqtype = 1  * codons
CodonFreq = 1  * F1X4
    clock = 0  * no clock
    model = 0  * one model for codons
  NSsites = 0  * one
    Mgene = 0  * rates
fix_kappa = 0  * kappa to be estimated
    kappa = 2  * initial kappa
fix_omega = 0  * estimate
    omega = .4 * initial omega
```

#### 1.7 Number of Rate Matrices

We varied the number $M$ (default 8) of rate matrices which determines the majority of parameters. Figure 6 shows that the accuracy is significantly smaller for $M = 2$ but does not vary greatly between $M = 3$ and $M = 12$. Even for the parameter-parsimonious $M = 3$ the AUC values are 0.991 (vertebrate), .967 (fly), .974 (yeast), when training on a mixture of clades and all other things being equal to the default version of ClaMSA ($M = 8$) whose values are in Table 1. These accuracies are not much worse than those of the default version of ClaMSA, suggesting that most of the gain is not due to the larger number of parameters of the model but likely rather to different training objective.

**Supplementary Table 1:** Accuracy Comparison. Best performances are in bold face.

| clade | method | AUC | errors | fpr at tpr 0.8 |
|---|---|---|---|---|
| vertebrate | ClaMSA | 0.992 | 68 | 0.3% |
| | ClaMSA vert | **0.992** | **64** | **0.2%** |
| | ClaMSA fly | 0.991 | 108 | **0.2%** |
| | ClaMSA yeast | 0.891 | 752 | 14.4% |
| | ClaMSA LogReg | 0.988 | 83 | 0.3% |
| | PhyloCSF | 0.945 | 447 | 2.6% |
| | codeml | 0.915 | 1021 | 8.7% |
| fly | ClaMSA | 0.971 | 236 | 3.2% |
| | ClaMSA vert | 0.950 | 321 | 4.4% |
| | ClaMSA fly | **0.979** | **195** | **1.8%** |
| | ClaMSA yeast | 0.795 | 842 | 28.3% |
| | ClaMSA LogReg | 0.961 | 291 | 4.4% |
| | PhyloCSF | 0.905 | 592 | 11.7% |
| | codeml | 0.877 | 957 | 15.2% |
| yeast | ClaMSA | **0.979** | **48** | **2.7%** |
| | ClaMSA vert | 0.695 | 238 | 64.8% |
| | ClaMSA fly | 0.736 | 310 | 45.8% |
| | ClaMSA yeast | 0.946 | 75 | 4.9% |
| | ClaMSA LogReg | 0.943 | 129 | 8.7% |
| | PhyloCSF | 0.956 | 356 | **2.7%** |
| | codeml | 0.812 | 440 | 32.1% |

#### 1.8 Accuracy Results

Table 1 shows the accuracy results for all three examined clades.

#### 1.9 Application Example

ClaMSA was run on all 12-way vertebrate alignments with an exon candidate on human chromosome 1. Hits which overlapped a known gene or repeat were filtered out. The example from Figure 7 was hand-picked. It shows a region, where ClaMSA identified with high confidence a coding region, but the UCSC browser database did not contain a coding gene entry.

Tyner, C., Barber, G. P., Casper, J., Clawson, H., Diekhans, M., Eisenhart, C., Fischer, C. M., Gibson, D., Gonzalez, J. N., Guruvadoo, L., Haeussler, M., Heitner, S., Hinrichs, A. S., Karolchik, D., Lee, B. T., Lee,

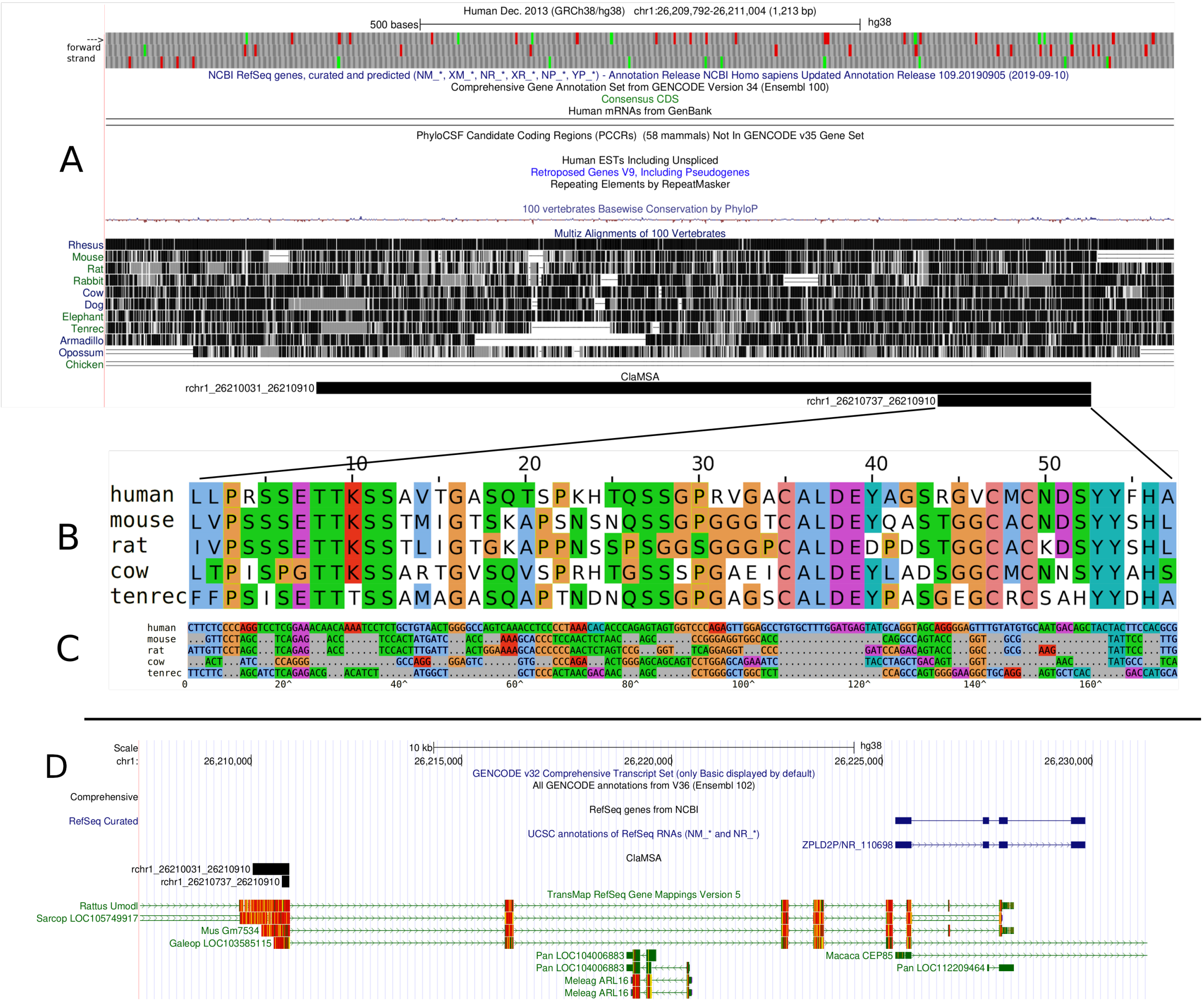

**Supplementary Figure 7:** Extended version of Figure 3 of the main document. **A**: UCSC genome browser. The genomes of human, mouse, rat, cow and tenrec contain contain longer ORFs of lengths 339, 619, 507, 386 and 360 amino acids length upstream of the respective genome positions aligned to the right boundary of the two ClaMSA regions. **B**: Amino acid alignment. **C**: Codon alignment. Alignments were viewed using `alv` (github.com/arvestad/alv). **D**: General human locus around discovery. The ClaMSA candidate coding regions (black bars to left) appear to be at the 5'-end of a human gene ZPLD2P with significantly truncated annotation, as additionally evidenced by a TransMap (Stanke et al., 2008) alignments of mouse gene Zpld2 and others.

C. M., Nejad, P., Raney, B. J., Rosenbloom, K. R., Speir, M. L., Villarreal, C., Vivian, J., Zweig, A. S., Haussler, D., Kuhn, R. M., and Kent, W. J. (2016). The UCSC Genome Browser database: 2017 update. *Nucleic Acids Research*, 45(D1):D626–D634.
